# Investigation on the Effectiveness of *Lactobacillus plantarum* in Enhancing the Folate content of *Injera* made with Different Cereals

**DOI:** 10.1101/2022.12.09.519824

**Authors:** Aynadis Tamene, Kaleab Baye, Tesfaye Mekuriyaw

## Abstract

Injera is an Ethiopian fermented pancake-like flatbread made from different cereals. The aim of the study was to investigate the effect of injera making process using different cereals (tef, sorghum, wheat and barley) on folate content and to evaluate the effectiveness of Lactobacillus plantarum (L. plantarum) in enhancing folate of injera made with different cereals. Cereals were used alone or in combination (tef and sorghum (1:1), wheat and sorghum (3:1), sorghum (100 %) and Barley (100 %). L. plantarum previously isolated from tef dough and ersho were used as starters. Folate content of flour, dough and injera was determined by microbiological assay. The contribution of consumption of injera made with different cereals to the folate requirement of children and women of reproductive age was evaluated. Acceptability of injera was estimated by 30 adult healthy volunteers using a 9-point hedonic scale. Among the studied cereals, the highest average folate content (49.9 μg/100 g) was observed in 100 % sorghum flour and the least (32.2 μg/100 g) in 100 % barley flour, on dry weight basis. After fermentation, the highest average folate content (60.1 μg/100 g) was observed in 100 % sorghum dough fermented with L. plantarum and the least (27.6 μg/100 g) was observed in the same dough but fermented with ersho. The average folate contents (fresh weight basis) of 100 % sorghum, wheat & sorghum (3:1), tef & sorghum (1:1) and 100 % barley injeras fermented with L. plantarum were 14.48 μg/100 g, 15.45 μg/100 g, 13.23 μg/100 g and 13.13 μg/100 g, respectively. Consumption of L. plantarum fermented injera made with different cereals can contribute up to 8 % of the recommended folate intake of women of reproductive age. Injera made with tef and sorghum blend (1:1) and fermented with L. plantarum was highly accepted.

## 1. Introduction

Folate (vitamin B9), plays a key role as an essential coenzyme in organisms to provide one-carbon units for nucleotide biosynthesis, amino acid metabolism, and DNA methylation (Crittenden *et al*., 2003; Iyer and Tomar, 2009; Rossi *et al*., 2011). Folate is a crucial component in maintaining the individual’s health over the course of life (Iyer & Tomar, 2009). Folate is also needed to prevent neural tube defects (NTDs) in the developing fetus. Inadequate intake of folate leads to folate deficiency (Hermann and Obeid, 2011a).

Folate deficiency is very frequent in many countries worldwide (McLean *et al*., 2008; Youngblood *et al*., 2013). In Africa, folate deficiency is mainly related to poor dietary diversity and reduced food intake (Dewey & Brown, 2003). The deficiency of folate has been implicated in a wide variety of disorders such as Alzheimer’s, coronary heart diseases, osteoporosis, increased risk of breast and colorectal cancer, poor cognitive performance, hearing loss and NTDs (LeBlanc, *et al*., 2007; LeBlanc *et al*., 2011). Inadequate maternal folate status has also been associated with low infant birth weight, preterm delivery and fetal growth retardation (Scholl & Johnson, 2000). Megaloblastic anemia and elevated blood concentrations of homocysteine have also been linked to folate deficiencies (Aslinia *et al*., 2006) and (Ho *et al*., 2011). Insufficient intake of folate is associated with high blood homocysteine concentration, a potential risk factor for coronary heart disease and stroke (Djuric *et al*., 2008). The gap between recommended and actual folate intake has led to mandatory folic acid fortification in more than 60 countries around the world. On the other hand, very high folic acid intake has been questioned, as it could mask vitamin B12 deficiency or promote the growth of pre-neoplastic lesions (Hermann and Obeid, 2011b) (Hoekstra *et al*., 2008).

Though most animal origin foods and some vegetables like dark-green leafy vegetables are rich sources of folate, regular consumption of these foods is limited in low and middle income countries (LMIC) (Zerfu *et al*., 2016). In addition, limited availability and access to folic acid fortified foods and low compliance/adherence to folic acid supplementation during pregnancy is prevalent (Taye *et al*., 2015). These combined effects lead a significant proportion of the population leaving in LMIC to be at a greater risk of folate deficiency and its adverse effects (McLean *et al*., 2008; Haidar, 2010).

Supplementation, food fortification, and dietary diversification are considered as main strategies to overcome micronutrient deficiencies (Bhutta *et al*., 2013). Food fortification and clinical trials consistently show that increasing woman’s intakes of folic acid during the periconceptional period results in a decrease in the prevalence of NTDs (Castillo-Lancellotti *et al*., 2013). Daily folic acid supplementation in pregnant women is recommended to reduce the risk of low birth weight, maternal anemia and NTD (WHO, 2012). Therefore, many countries have adopted the mandatory fortification of staple foods with folic acid to maximise the proportion of women of childbearing age having optimal folate status. But contradicting research results are also reported on the negative health effect of the long term consumption of the synthetic folic acid which could increase the risk of developing autism (DeSoto *et al*., 2012).

These days a more natural and safe way of enhancing folate status of the population is emerging. For example, in recent years, natural folate enhancement in fermented food products by introducing folate producing microorganisms has gained more attention. *Lactic acid bacteria* (LAB) and yeasts which are the main actors of fermentation, could produce up to 100 μg/L and 145 μg/g cell mass, respectively (Sybesma *et al*., 2003; Hjortmo *et al*., 2005; Iyer and Tomar, 2009). Total folate production up to 256 μg /L by genetically engineered *L. lactis ssp. lactis* was also reported by Sybesma *et al*., (2003).

Cereal-based foods are widely consumed as the primary staple foods in many countries of the world, especially in Africa (Tamene *et al*., 2019a). Many cereal-based staple foods are usually undergoing a fermentation step before consumption (Humblot & Guyot, 2008). For example, *Injera* is a leavened, flat and round cereal-based fermented bread made from different cereals (Mengesha *et al*., 2022). It is a staple food that is widely consumed in Ethiopia (Baye *et al*., 2013).

As *Injera* made with tef is widely consumed especially in Addis Ababa (capital city of Ethiopia), investigation on the effect of *Injera* processing (fermentation and baking) was performed on samples collected from Addis only (Tamene *et al*., 2019a). In another investigation, Tamene and his colleagues have also isolated a folate producing LAB (*L. plantarum*) from tef fermentation and the bacteria was found to be successful in enhancing the folate content of *Injera* made with tef (Tamene *et al*., 2019b; Tamene *et al*., 2022).

Different factors may affect the ability of *L. plantarum* on folate production during fermentation which may include the raw materials (different cereals) which could be used for *injera* preparation. The method of processing could also affect folate content of the food. In Ethiopia, research has been conducted to investigate the effect of *Injera* preparation on folate concentration. However, emphasis was given only on *Injera* made with tef. Therefore, this research aimed to further investigate the effect of *injera* preparation made with different cereals on folate content. In addition, it investigated the possibility of enhancing folate content of *injera* made with different cereals using *L. plantarum* previously isolated from tef fermentation as a starter.

## 2. Material and method

### 2.1. Chemicals and raw materials

Unless otherwise specified, all the necessary chemicals and reagents used in this work were purchased from Sigma-Aldrich Chemie GmbH (Buchs, Switzerland). The raw material (tef, wheat, sorghum and barley) were obtained from local market of Addis.

### 2.2. Sampling and sample preparation

Information on the different types of cereals used for *injera* making was obtained from the literature. The type of cereals and the most common blend proportions used in the preparation of *injera* were taken from Baye *et al*., (2013) and from actual observation from selected households. According to Baye *et al*., (2013), tef and white sorghum were consumed in the lowland and barley, wheat and red sorghum were consumed in highlands of Ethiopia.

Accordingly, cereals such as sorghum, tef, wheat and barley were selected for the current study. The blending ratio of cereal types to make *injera* was noted as sorghum (100 %), sorghum & tef (1:1), wheat & sorghum (3:1) and barley (100 %) (table 1). Samples of different cereals were collected from the local markets of Addis Ababa. From each cereal types, 5kg sample were bought and transported to Addis Ababa University, Center for Food Science and Nutrition laboratory for laboratory analysis. The different cereal types have different preparation steps; Tef: cleaned, sieved and milled, Sorghum: cleaned, decorticated and husk removed and milled, Wheat: cleaned and milled and Barley: cleaned, conditioned, husk removed and milled. The flour for different cereal types passed through a 0.5 mm sieve. Flour were blended using dry particulate blender and subjected into four experimental runs (table 1).

**Table 1.** Blend of cereal flour used for the preparation of *injera*

| Experimental Run | Blends of cereal flour |
| --- | --- |
| 1 | Tef : Sorghum (1:1) |
| 2 | Wheat : Sorghum (3:1) |
| 3 | Sorghum (100 %) |
| 4 | Barley (100 %) |

### 2.3. Preparation of *Injera* made with *L. plantarum* and *ersho* as innoculums

#### 2.3.1. Preparation of the inoculums

Two different inoculums were prepared. A leftover from previous successful spontaneous fermentation batch (*ersho*) which was taken from household was used as inoculum to prepare traditional *injera*.

The folate producing *L. plantarum P2R3FA*, previously isolated from fermented tef batter (Tamene *et al*., 2019a) was cultivated by streaking the strain conserved at -80 C in De Man, Rogosa, and Sharpe (MRS) broth and glycerol (40%) on MRS agar and incubated at 30 °C for 48 h. A colony was collected from each pure culture plate, grown in MRS broth (24 h, 30°C), and centrifuged (14,000 x g, 7 min). The pellets were washed twice with the same volume (9 mL) of sterile saline solution (0.9 % NaCl) and re-suspended in the same volume of solution. The final suspension contained around 10^9^ colony-forming units (cfu)/mL. The inoculum for *injera* fermented with the folate-producing *L. plantarum P2R3FA* was prepared by mixing this suspension with flour (1:1) (v/w).

#### 2.3.2. Preparation of *Injera* made with different cereals and fermented with the above indicated inoculums

The traditional *injera* making process was developed after detail observations from 10 selected households. Household were selected deliberately (where blended cereals are used for *injera* making) and resulted in a flow diagram presented in **fig 1**. Briefly, dough was prepared by using blended and/or whole flour of (0.5 kg), 0.75 L sterile tap water and 125 mL starter culture (0.5:0.75:0.125) (w/v/v). The 125 mL starter culture was prepared by mixing 62.5 mL saline containing the *L. plantarum* strain and 62.5 g flour (1:1) (v/g), similar proportion was also used for *ersho* starter. Thereafter, 200 mL sterile tap water was carefully added to cover the surface of the batter and incubated for 4 days at room temperature (1st phase fermentation). After 4 days of fermentation, liquid present on top of the batter was decanted and replaced by a similar amount of fresh tap water except for barley fermentation where there was no supernatant. Before baking *injera* from the fermented batter, 1/11th of the fermented batter was mixed with sterile tap water (1:3) (v/v), boiled for about 10 min and cooled to a temperature around 45 °C. The resulting product (*absit*) was added back to the remaining fermented batter to enhance proper fermentation. Incase of barley fermentation, sterile cold water was used instead of *absit*. The batter was allowed to ferment for 2 h at 25 °C until gas production was observed (2^nd^ phase fermentation). Finally, the fermented liquid batter (450 mL) was poured onto a hot clay, enclosed and baked for 2 min. The final flat pancake-like product is known as *injera*. Three bakings were performed for each inoculum mentioned under section 2.3.1.

**Fig 1.**
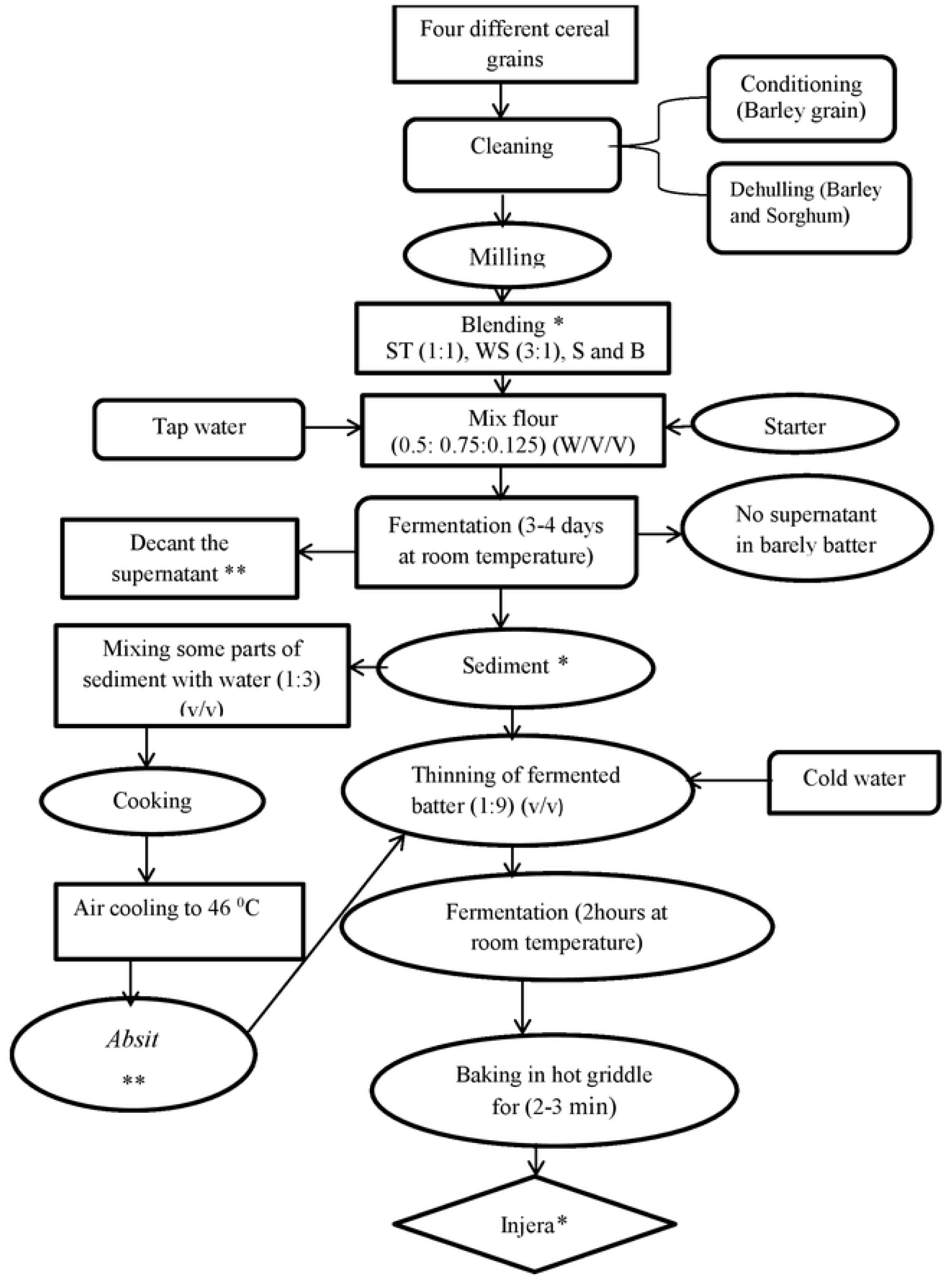
Description of processing of *injera made with different cereal blends and using* different starters. ST= sorghum & tef flour (1:1), WS= wheat & sorghum flour (3:1), S= sorghum flour (100 %) and B= barley flour (100 %). * indicates sample were taken for folate analysis ** indicates no *absit* preparation and decantation process during barely fermentation. **Starters:** *Ersho* (traditional backslopping) Folate producing *L. plantarum* P2R3FA

#### 2.3.3. Sampling

During *injera* preparation, pH of the batter was measured initially before fermentation and after second stage fermentation. Fermented batter was sampled to measure dry matter (DM) and folate contents before the addition of *absit*, and *injera* was also sampled to measure DM and folate contents and subjected to sensory analysis. DM was analyzed by oven drying at 105 °C. pH was measured using an aliquot of dough immediately after diluting it with deionized water (1:1) (v/v).

### 2.4. Folate measurement and effect of processing

#### 2.4.1. Folate measurement

The total folate content of tef flour, dough and *injera* was determined in triplicates using the reference microbiological method, after trienzyme extraction (Kariluoto *et al*., 2004; Tamene *et al*., 2019b). The total folate content was determined based on the growth of folate dependent strain, *Lactobacillus rhamnosus* ATCC 7469 as the test organism and (6S)-5-formyltetrahydrofolate (5-HCO-H4 folate) as the calibrant. Performance of the method was checked by testing a blank sample and certified reference material (BCR-121 whole meal flour). Only folate contents in the range of certified value (500 ± 70 ng/g dry matter) were accepted. In addition, for triplicate samples, folate content variations >10 % were not accepted. Analytical procedures were carried out under yellow or subdued light. Alternatively, samples and calibrants were covered with aluminum foil. Sample extracts were kept under nitrogen atmosphere whenever necessary.

#### 2.4.2. Effect of processing

The effect of cooking (baking), fermentation and the overall *injera* processing on the total folate content was evaluated and results were expressed as percentage retention.

### 2.5. Contribution of *injera* consumption to the recommended nutrient intake (RNI) of folate

Based on the existing data on *injera* consumption from the Ethiopian National Food Consumption survey (EPHI, 2013) and RNI of folate from world health organization (FAO/WHO, 2004), the contribution of consumption of *injera* made with different cereals to RNI of folate was estimated for children and women of reproductive age. This population groups were selected because they represent a higher risk for folate deficiency.

### 2.6. Sensory evaluation

The aim of this experiment was to determine whether the folate-enriched *injera* made with different cereals are acceptable or not by potential consumers. The acceptability and sensory profile for the prepared *injera* was estimated by 30 adult healthy volunteers (women of reproductive age), using a 9-point hedonic scale. The thirty judges of adult healthy volunteers were selected at random on March, 2022. They were provided with the randomly sequenced samples for testing. They were asked to evaluate the products for color, taste, odor, texture (mouth feel), and overall acceptability. The judges used a 9-point hedonic scale where 1 = liked extremely and 9 = disliked extremely to evaluate the acceptability of *injera*. They were instructed to cleanse their mouth with water before testing the next sample. Informed consent was obtained from all participants in the study and ethical approval was obtained from the Institutional Review Board of the College of Natural and Computational Sciences of Addis Ababa University. All authors had access to all relevant information to identify study participants both during and after data collection.

### 2.7. Statistical analysis

Statistical analysis of folate and sensory acceptability was computed using SPSS version 25 software. The folate analyses were carried out in triplicate and the average values and standard deviations were calculated. The differences between means of folate values in different cereal flour, dough and *injera* and sensory acceptability were quantified using analysis of variance (ANOVA) and Tukey’s post hoc test was used to compare the mean values. Significant mean differences were considered with at *P* value ≤ 0.05.

## 3. Result

### 3.1 pH of the dough

The pH of initial dough made with four different blended cereals recorded at 25 ºC ranged from 5.5 to 6.1 with an average of 5.8 ± 0.3. After fermentation, the pH value of doughs made with different cereals and fermented with *L. plantarum* ranged from 3.38 to 3.43 with an average value of 3.36 ± 0.07. Whereas the pH value of *ersho* fermented doughs made with different cereals ranged from 3.30 to 3.42 with an average value of 3.41 ± 0.09. There was no significant difference on pH value of all doughs regardless of the cereal types and starters used.

### 3.2 DM content of flour, dough and *injera*

The DM content of flour, dough and *injera* made with different cereals and starters were analyzed and results are shown in **Table 2**. The average DM of different flour was ranged from 91.7 ± 0.57 to 92.7 ± 0.57 % by mass. There was no significant difference on DM content of all the flours.

**Table 2.** DM content of flour, dough and *injera (% by mass)*

| Blends | Flour | Dough fermented with: |  | *Injera* fermented with: |  |
| --- | --- | --- | --- | --- | --- |
|  |  | *Ersho* | *L. plantarum* | *Ersho* | *L. plantarum* |
| Tef and sorghum (1:1) | $91.7 \pm 0.57^a$ | $39.8 \pm 0.15^b$ | $38.6 \pm 0.43^a$ | $45.3 \pm 0.26^b$ | $43.5 \pm 0.3^b$ |
| Wheat and sorghum (3:1) | $91.7 \pm 0.57^a$ | $39 \pm 0.93^b$ | $39.5 \pm 0.35^a$ | $41.9 \pm 0.53^a$ | $43.5 \pm 0.2^b$ |
| Sorghum (100 %) | $92.7 \pm 0.57^a$ | $40.8 \pm 0.25^b$ | $41.5 \pm 0.35^b$ | $38.2 \pm 0.76^a$ | $44.7 \pm 0.5^b$ |
| Barley (100 %) | $92.7 \pm 0.57^a$ | $35.6 \pm 0.31^a$ | $37.9 \pm 0.94^a$ | $37.2 \pm 0.53^a$ | $39.9 \pm 0.4^a$ |

Values in the table are mean and standard deviation of triplicate samples.
Different superscript letters along the column indicates a statistically significant difference at ( $p < 0.05$ )

The average DM contents of *ersho* fermented dough ranged from 35.6 ± 0.31 to 40.8 ± 0.25 % by mass. In case of *L. plantarum* fermented dough, the DM was in the range of 37.9 ± 0.94 and 41.5 ± 0.35 % by mass. The result has showed that significantly higher DM (41.5 ± 0.35 % by mass) was observed on dough made with sorghum whereas there was no significant difference on DM content between other doughs.

The average DM content of *injera* made with *ersho* ranged from 37.2 ± 0.53 to 45.3 ± 0.26 % by mass. With the exception of *injera* made with tef and sorghum blend, which had significantly higher DM (45.3 ± 0.26), there was no significant difference on DM contents of all *injera* fermented with *ersho*. Regarding *injera* fermented with *L. Plantarum*, the average DM content ranged from 39.9 ± 0.4 to 44.7 ± 0.5 % by mass. Our result indicated that significantly lower DM (39.9 ± 0.4 % by mass) was observed on *injera* made with barley otherwise there was no significant difference on DM content between other *injeras*.

### 3.3 Folate content of flour, dough and *injera* made with different cereals and fermented with different starters

The folate contents of flour, dough and *injera* made with different cereals and fermented with different starters were analyzed and results are shown in **Table 3**. The average folate contents of different blends of flour were ranged from 32.2 ± 1.5 to 49.9 ± 1.1 μg/100g. Significantly higher folate content (49.9 ± 1.1 μg/100g) was recorded in sorghum flour whereas significantly lower folate content (32.2 ± 1.5 μg/100g) was observed in barley flour. There was no significant difference between average folate content of tef and sorghum (1:1) and wheat and sorghum (3:1) blended flour, which had 42.6 ± 1.4 and 40.4 ± 1.5 μg/100g of folate, respectively.

**Table 3.** Folate content (μg/100g) of flour, dough and *injera* fermented with *L. plantarum* and ersho, DM basis

| Blends | Flour | Dough fermented with: |  | *Injera* fermented with: |  |
| --- | --- | --- | --- | --- | --- |
|  |  | *Ersho* | *L.plantarum* | *Ersho* | *L.plantarum* |
| Tef and sorghum (1:1) | $42.6 \pm 1.4^b$ | $29.1 \pm 0.9^{aB}$ | $53.5 \pm 3.5^{bC}$ | $17.5 \pm 0.7^{aA}$ | $30.2 \pm 0.9^{aB}$ |
| Wheat and sorghum (3:1) | $40.4 \pm 1.5^b$ | $32.2 \pm 1.3^{bcB}$ | $56.9 \pm 1.6^{bcC}$ | $23.6 \pm 0.7^{bA}$ | $35.4 \pm 1.3^{abB}$ |
| Sorghum (100 %) | $49.9 \pm 1.1^c$ | $27.6 \pm 2.7^{aB}$ | $60.1 \pm 1.6^{cD}$ | $23.4 \pm 1.1^{bA}$ | $32.3 \pm 5.1^{aC}$ |
| Barley (100 %) | $32.2 \pm 1.5^a$ | $34.5 \pm 0.7^{cB}$ | $45.5 \pm 1.3^{aC}$ | $20.8 \pm 0.9^{bA}$ | $32.7 \pm 1.5^{aB}$ |

Values are means and standard deviations of triplicate samples.
Different lower case superscript letters along the column indicate a statistically significant difference at ( $p < 0.05$ ) and different upper case superscript letters across the row indicate a statistically significant difference at ( $p < 0.05$ )

The average folate contents of different dough fermented with *ersho* were ranged from 27.6 ± to 34.5 ± 0.7 μg/100g. The highest folate content (34.5 ± 0.7 μg/100g) was recorded in barley batter, however it was not statistically different from folate of wheat and sorghum blended dough (32.2 ± 1.3 μg/100g) at p ≥ 0.05. The lowest concentrations of folate in *ersho* fermentation was observed in sorghum batter (27.6 ± 2.7 μg/100g). However, it was not statistically different from folate of tef and sorghum blended dough (29.1 ± 0.9 μg/100g).

In case of *L. plantarum* strain fermented doughs, the average folate contents were in the range of 45.5 ± 1.3 and 60.1 ± 1.6 μg/100g. The highest folate content (60.1 ± 1.6 μg/100g) was recorded in sorghum batter, however it was not statistically different from folate of wheat and sorghum blended dough (56.9 ± 1.6 μg/100g) at (p ≥ 0.05). Significantly lower concentration of folate was observed in *L. plantarum* fermented barley batter (45.5 ± 1.3 μg/100g). Whereas, there was no significant difference between folate contents of wheat & sorghum (3:1) and sorghum & tef (1:1) blended doughs.

The average folate contents of *injera* made with different cereals and fermented using *ersho* were ranged from 17.5 ± 0.7 to 23.6 ± 0.7 μg/100g. Among the different blends, significantly lower folate contents (17.5 ± 0.7 μg/100g) was observed in tef and sorghum blended *injera*. However, there was no significant difference on folate contents among other *injera* samples at (p ≥ 0.05).

Results of the folate contents of *L. plantarum* fermented *injera* made with different cereals are also shown in **Table 3**. The average folate contents were in the range of 30.2 ± 0.9 and 35.4 ± 1.3 μg/100g. There was no significant difference on folate contents of all *injeras* fermented with *L. plantarum*. The highest (35.4 ± 1.3 μg/100g) and lowest (30.2 ± 0.9 μg/100g) folate contents were observed in wheat and sorghum and tef and sorghum blends, respectively.

The feasibility of *L. plantarum* on enhancing folate content of dough and *injera* made with different cereal blends were evaluated and compared with the folate contents of dough and *injera* fermented with *ersho*. In all cases, doughs fermented with the strain (*L. plantarum*) were found to have significantly higher folate contents as compared to all doughs fermented with *ersho* (**Table 3**). Looking at specific doughs, tef and sorghum (1:1) blended dough had 29.1 ± 0.9 and 53.5 ± 3.5 μg/100g, wheat and sorghum (3:1) blended dough had 32.2 ± 1.3 and 56.9 ± 1.6 μg/100g, sorghum dough had 27.6 ± 2.7 and 60.1 ± 1.6 μg/100g and Barley dough had 34.5 ± 0.7 and 45.5 ± 1.3 μg/100g folate contents when they were fermented with *ersho* and *L. plantarum* strain, respectively.

Similarly, in all cases, *injeras* fermented with our strain (*L. plantarum*) had significantly higher folate contents as compared to all *injeras* fermented with *ersho* as shown under **Table 3**. Looking at specific *injeras*, tef and sorghum (1:1) blended *injera* had 17.5 ± 0.7 and 30.2 ± 0.9 μg/100g, wheat and sorghum (3:1) blended *injera* had 23.6 ± 0.7 and 35.4 ± 1.3 μg/100g, sorghum *injera* had 23.4 ± 1.1 and 32.3 ± 5.1 μg/100g and Barley *injera* had 20.8 ± 0.9 and 32.7 ± 1.5 μg/100g folate contents when they were fermented with *ersho* and strain, respectively.

### 3.4 Effect of fermentation on folate content of dough

Percentage retention of folate after fermentation was calculated by comparing the amount of folate content in tef flour and in the dough on DM basis (**Fig 2**). A value above 100 % indicated that fermentation led to increases in folate whereas retention values less than 100 % indicated consumption/losses of folate.

**Fig 2.**
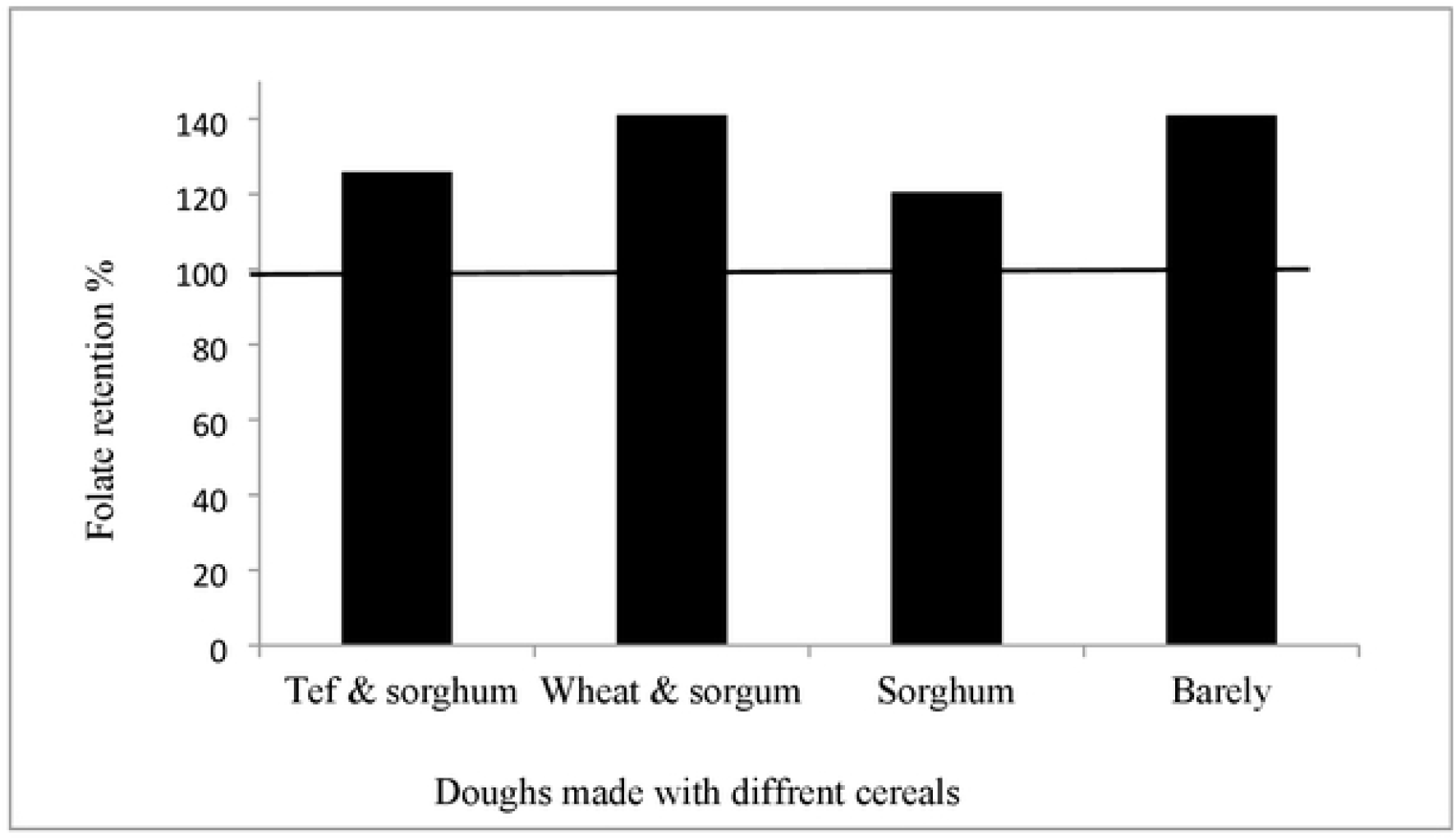
Folate retention (%) due to fermentation using *L. plantarum* Folate retention (%) = Folate dough / Folate flour x 100

Folate retention due to fermentation of different cereals fermented with *L. plantarum* ranged from 120 to 141 %. This showed more than 100 % folate retention for all of doughs fermented with the strain regardless of the blend used. The highest retention (141 %) was recorded in dough made with wheat & sorghum blend (3:1).

Folate retention due to fermentation of different cereals fermented with *ersho* ranged from 68 to 107 % (**Fig 3**). This showed less than 100 % folate retention for all of doughs with the exception of barley which had folate retention of 107 %. The higher retention for barley dough may be due to the absence of decantation process which may prevent the losses of folate and/or due to the presence of naturally occurring folate producing microorganisms involved in the fermentation process.

**Fig 3.**
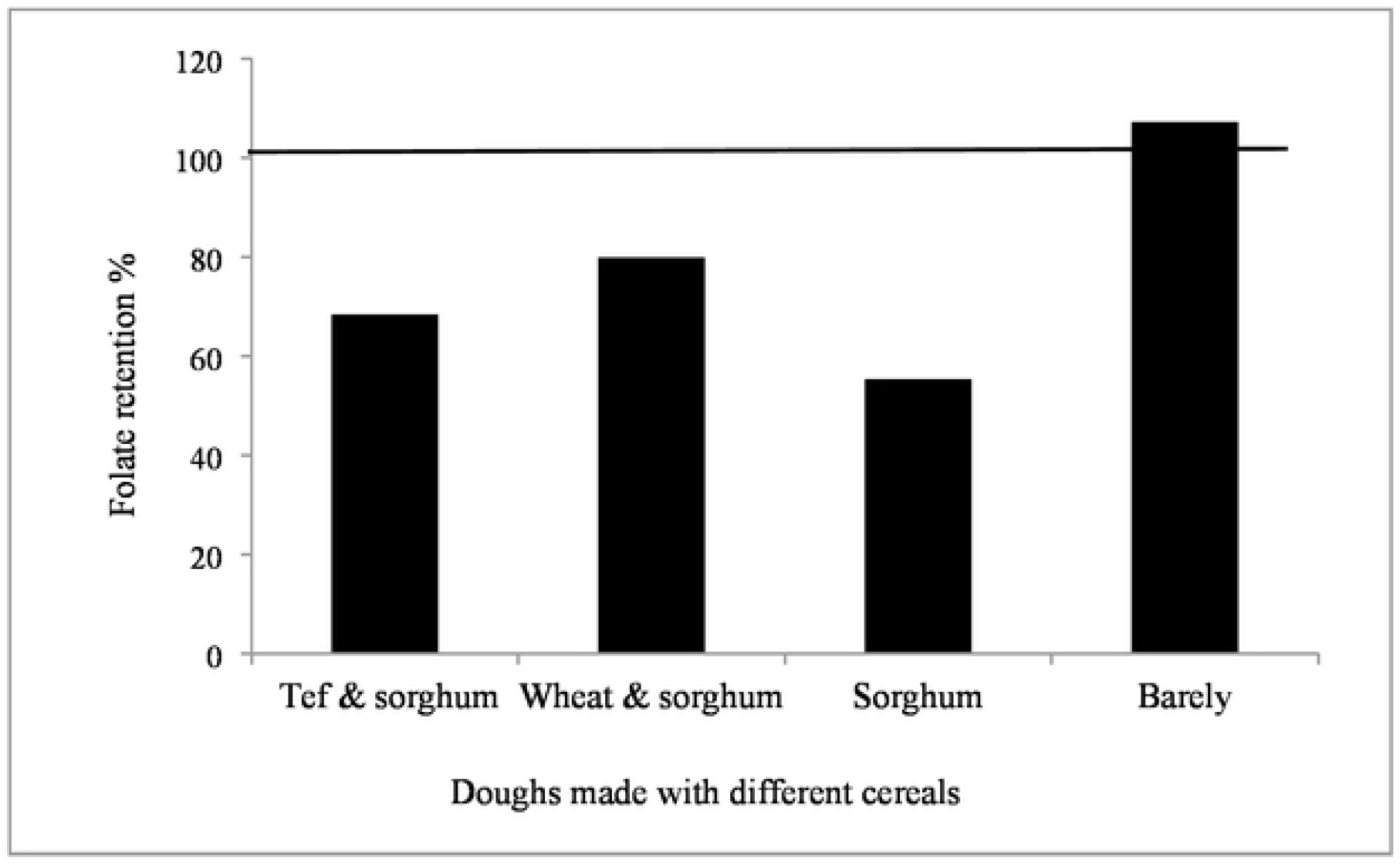
Folate retention (%) due to fermentation using *ersho* Folate retention (%) = Folate dough/Folate flour x 100

### 3.5 Effect of baking on folate content of *injera* made using *ersho* and *L. plantarum*

In all *L. plantarum* fermented *injera*s, regardless of the type of cereal blends, the folate retention value due to baking was less than 100 % and it ranged from 56 to 76 % with average folate retention of 66 % (**Fig 4**). The highest (76 %) and least (56 %) folate retention was observed in barely and sorghum *injera*s, respectively.

**Fig 4.**
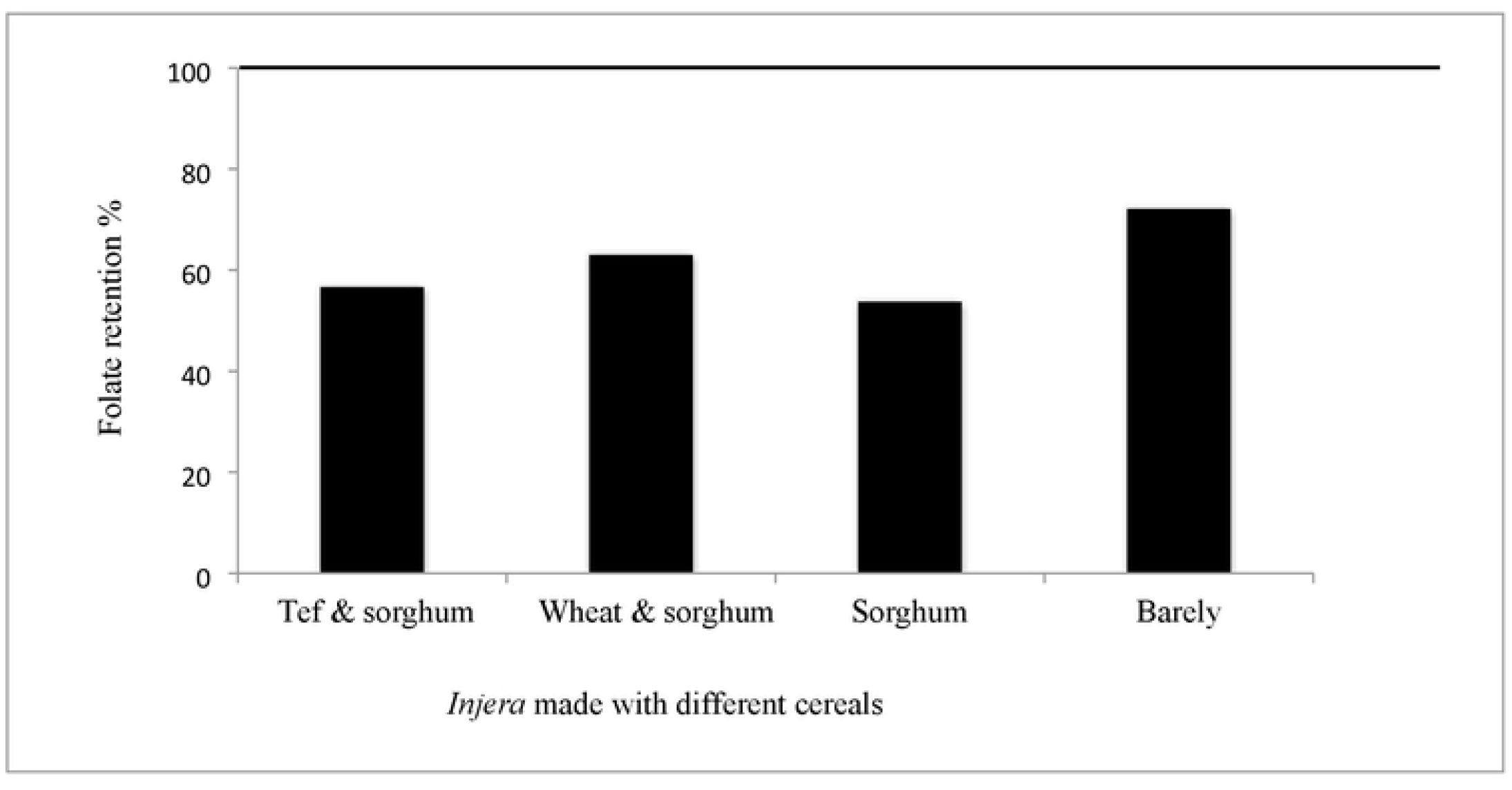
Folate retention (%) during baking of *injera* fermented using *L. plantarum* Folate retention (%) = Folate dough/Folate *injera* x 100.

Similarly, in all of *injeras* fermented with *ersho*, regardless of the type of cereal blends, the folate retention value due to baking was less than 100 % and it ranged from 60 to 76 % with average folate retention of 68 % (**Fig 5**). The highest folate retention (76 %) was observed in sorghum and the least folate retention (60 %) was recorded in barely.

**Fig 5.**
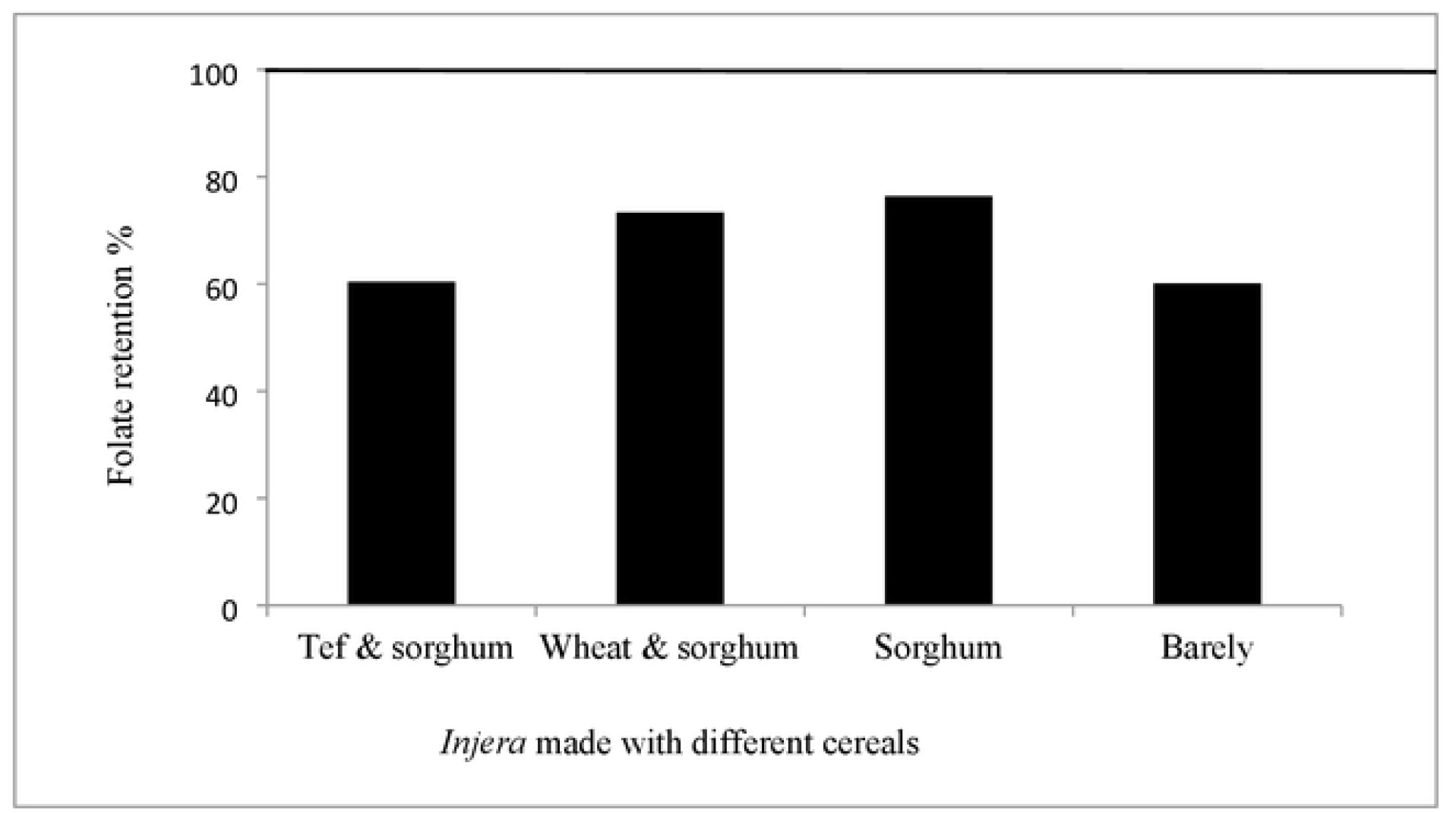
Folate retention (%) during baking of *injera* fermented using *ersho* Folate retention (%) = Folate dough/Folate *injera* x 100.

### 3.6 Folate content and calculated contribution of *injera* made with different cereals and fermented with different starters to folate requirements (RNI)

The average folate content of *injera* made with different cereals and fermented with *ersho* was in the range of 7.82 ± 0.61 and 9.79 ± 0.28 μg/100 g per fresh weight (FW). The lowest average folate content (7.82 ± 0.61 μg/100 g FW) was recorded in barley *injera* and the highest (9.79 ± 0.28 μg/100 g FW) was observed in wheat and sorghum blend (3:1) *injera* **(Table 4**). Whereas, the average total folate content of *injera* made with different cereals and fermented with *L. plantarum* was ranged from 13.13 ± 0.59 to 15.45 ± 0.58 μg/100 g FW. The least folate content (13.13 ± 0.59 μg/100 g FW) was observed in barley and the highest (15.45 ± 0.58 μg/100 g FW) was noted in wheat and sorghum blend (3:1) *injera*.

**Table 4.** Folate content and calculated contribution of *injera* made with different cereal blends and fermented with *ersho* and *L. plantarum* to folate intakes requirements

| Starters used | *Injera* | Folate<br>($\mu\text{g}/100$ g FM) | Contribution to RNI (%) | |
| --- | --- | --- | --- | --- |
|  |  |  | Children<br>(1-3 years) | Women (19 years and older) |
| *Erscho* | Tef & sorghum (1:1) | $7.99 \pm 0.34$ | 3.5 | 4.1 |
| | Wheat & sorghum (3:1) | $9.79 \pm 0.28$ | 4.3 | 4.9 |
| | Sorghum | $8.94 \pm 0.74$ | 3.9 | 4.5 |
| | Barley | $7.82 \pm 0.61$ | 3.4 | 3.9 |
| *L. plantarum* | Tef & sorghum (1:1) | $13.23 \pm 0.43$ | 5.8 | 6.7 |
| | Wheat & sorghum (3:1) | $15.45 \pm 0.58$ | 6.8 | 7.8 |
| | Sorghum | $14.48 \pm 2.30$ | 6.3 | 7.3 |
| | Barley | $13.13 \pm 0.59$ | 5.7 | 6.6 |

Values of folate are means ± standard deviation
Average *injera* consumption by children = 66 g/day, and by women of reproductive age = 202 g/day (Tamene *et al.*, 2019a).

Contribution of *injera* consumption to folate requirements was calculated using current level of average *injera* consumption (66 g/day) for children and (202 g/day) for women of reproductive age (Tamene *et al*., 2019a) and RNI of folate (150 μg/day) for children and (400 μg/day) for women of reproductive age (WHO, 2004). The least folate contribution to RNI was observed in *injera* fermented by traditional process with *ersho* and the contribution was in the range of 3.4 to 4.3 % for children and 3.9 to 4.9 % to women of reproductive age. The highest contributions to the folate RNI to both population group was observed by wheat and sorghum *injera* and which contributed 4.3 % and 4.9 % to children and women of reproductive age, respectively. However, the least contribution to folate RNI to both population group was observed in barley *injera*, which contributed 3.4 % and 3.9 % to children and women of reproductive age, respectively.

With regard to *L. plantarum* fermented *injera*, the folate contributions to RNI was in the range of 5.7 to 6.8 % and 6.6 to 7.8 % to children and women of reproductive age, respectively. The highest contributions for both population groups was observed in wheat and sorghum *injera* (3:1), which contributed 6.8 % and 7.8 % to children and women of reproductive age, respectively. However, the least contribution to folate RNI for both population groups was observed in barley *injera* with 5.7 % and 6.6 % to children and women of reproductive age, respectively.

### 3.7 Sensory acceptability of different *injera* made using *L. plantarum* and *ersho*

As per the panellists, all of the samples made using different cereals and selected inoculums were found to be in the acceptable range with the exception of sorghum *injera*, which was rated as neither liked nor disliked by the panellists **(Table 5)**. The overall acceptability test result showed that sorghum *injera* fermented with both inoculums (*ersho* and *L. plantarum* P2R3FA) was the least acceptable product whereas tef and sorghum blended and 100 % barley *injeras* fermented with both inoculums were the most acceptable products. Looking at each specific parameter, with the exception of colour, for all other sensory parameters (taste, odour, texture and appearance), wheat and sorghum blended and 100 % sorghum *injeras* fermented with both inoculums were found to be the least liked product.

**Table 5.**
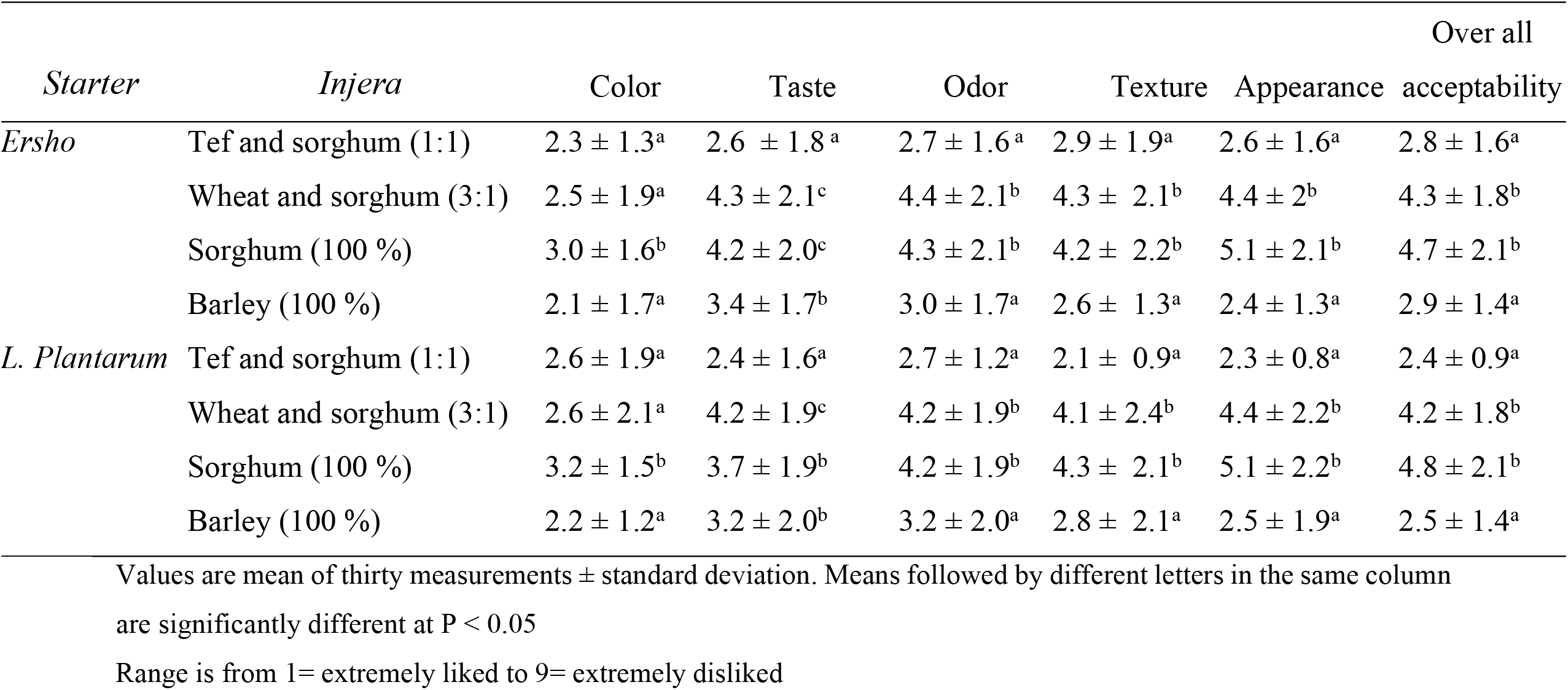
Sensory acceptability of different *injera* fermented with *ersho* and *L. plantarum* starters

| Starter | Injera | Color | Taste | Odor | Texture | Appearance | Over all |
| --- | --- | --- | --- | --- | --- | --- | --- |
|  |  |  |  |  |  |  | acceptability |
| *Erscho* | Tef and sorghum (1:1) | 2.3 ± 1.3<sup>a</sup> | 2.6 ± 1.8<sup>a</sup> | 2.7 ± 1.6<sup>a</sup> | 2.9 ± 1.9<sup>a</sup> | 2.6 ± 1.6<sup>a</sup> | 2.8 ± 1.6<sup>a</sup> |
|  | Wheat and sorghum (3:1) | 2.5 ± 1.9<sup>a</sup> | 4.3 ± 2.1<sup>c</sup> | 4.4 ± 2.1<sup>b</sup> | 4.3 ± 2.1<sup>b</sup> | 4.4 ± 2<sup>b</sup> | 4.3 ± 1.8<sup>b</sup> |
|  | Sorghum (100 %) | 3.0 ± 1.6<sup>b</sup> | 4.2 ± 2.0<sup>c</sup> | 4.3 ± 2.1<sup>b</sup> | 4.2 ± 2.2<sup>b</sup> | 5.1 ± 2.1<sup>b</sup> | 4.7 ± 2.1<sup>b</sup> |
|  | Barley (100 %) | 2.1 ± 1.7<sup>a</sup> | 3.4 ± 1.7<sup>b</sup> | 3.0 ± 1.7<sup>a</sup> | 2.6 ± 1.3<sup>a</sup> | 2.4 ± 1.3<sup>a</sup> | 2.9 ± 1.4<sup>a</sup> |
| *L. Plantarum* | Tef and sorghum (1:1) | 2.6 ± 1.9<sup>a</sup> | 2.4 ± 1.6<sup>a</sup> | 2.7 ± 1.2<sup>a</sup> | 2.1 ± 0.9<sup>a</sup> | 2.3 ± 0.8<sup>a</sup> | 2.4 ± 0.9<sup>a</sup> |
|  | Wheat and sorghum (3:1) | 2.6 ± 2.1<sup>a</sup> | 4.2 ± 1.9<sup>c</sup> | 4.2 ± 1.9<sup>b</sup> | 4.1 ± 2.4<sup>b</sup> | 4.4 ± 2.2<sup>b</sup> | 4.2 ± 1.8<sup>b</sup> |
|  | Sorghum (100 %) | 3.2 ± 1.5<sup>b</sup> | 3.7 ± 1.9<sup>b</sup> | 4.2 ± 1.9<sup>b</sup> | 4.3 ± 2.1<sup>b</sup> | 5.1 ± 2.2<sup>b</sup> | 4.8 ± 2.1<sup>b</sup> |
|  | Barley (100 %) | 2.2 ± 1.2<sup>a</sup> | 3.2 ± 2.0<sup>b</sup> | 3.2 ± 2.0<sup>a</sup> | 2.8 ± 2.1<sup>a</sup> | 2.5 ± 1.9<sup>a</sup> | 2.5 ± 1.4<sup>a</sup> |

Values are mean of thirty measurements ± standard deviation. Means followed by different letters in the same column
are significantly different at P < 0.05
Range is from 1= extremely liked to 9= extremely disliked

## 4. Discussions

In Ethiopia, *injera* is mainly produced from tef, but it could be also prepared from other different cereals like barley, sorghum and wheat either alone or in combination. Few studies focused on investigating the effect of tef *injera* preparation on folate contents and an attempt was done to enhance the folate content of tef *injera* through the application of folate producing bacteria (*L. plantarum*), previously isolated from tef fermentation. Therefore, the current study was conducted to investigate the effect of *injera* preparation made with different cereals on folate content. The effectiveness of the previously isolated *L. plantarum* in improving the folate content of *injera* prepared using other cereals like wheat & sorghum blend (3:1), tef & sorghum blend (1:1), sorghum (100 %) and barley (100 %) was investigated. The contribution of *injera* made with different cereal types and fermented with *ersho* and *L. plantarum* to folate RNI was investigated. The study also focused on evaluating sensory profile of *injera* made with different cereals and fermented with *ersho* and *L. plantarum*.

The current study shows that all of the cereals flour used in our study are relatively good source of folate. The impact of fermentation (one of the steps in *injera* preparation) was hugely variable. In the case of doughs fermented with *L. plantarum*, though highly variable, fermentation leads to net production of folate. Whereas, those doughs fermented with the traditional process using *ersho* leads to reduction of folate with the exception of barley dough which may be due to the presence of other naturally occurring folate producing microorganisms and lack of decantation process which may lead to folate loses. In contrast with fermentation, baking invariably led to folate losses. The current finding is exactly in agreement with a study report by Tamene *et al*. (2019a), where they reported that tef fermentation could either increase or decrease folate contents but baking always leads to loses.

Though slightly lower, the average total folate content of flours of the different cereals used in this stud (49.9 μg/100 g DM in sorghum, 42.6 μg/100 g DM in tef and sorghum blend (1:1), 40.4 μg/100 g DM in wheat and sorghum blend (3:1), are comparable with previously reported folate of tef flour (52.1 μg/100 g DM) (Tamene *et al*., 2019a). Folate of barley flour was lower than flour of tef. It was also lower than all other reported folate values of other cereals including oats, rice, whole wheat flour and maize (30–40 μg/100 g) and sorghum (77.0 μg/100 g) (Hager *et al*., 2012).

As indicated by current studies, fermenting food products like cereals and dairy products has the potential to enhance folate content, and its effectiveness as a strategy to combat health complications of folate deficiencies needs to be further investigated (Rollán *et al*., 2019; Saubade *et al*., 2017). Our study has also revealed the possibility of increasing folate by fermentation as more than 100 % folate retention was observed in barley fermentation. This result suggests that folate producing microorganisms may dominate those that do not produce or consume folate as previous studies confirmed the co-occurrence of both folate producing and consuming microorganisms during fermentation process (Okoroafor *et al*., 2019). The type of microorganisms and the overall conditions that led to folate production in barley fermentation needs further detail examination. Fermentation of all cereals using *L. plantarum* always leads to net folate production, which showed that this strain is as effective as it was in tef fermentation in enhancing the folate content.

As baking is the second major step in *injera* preparation, any folate that is originally present in a food or produced as a result of fermentation need to survive the baking temperature. Folate is known to be heat sensitive and processing and cooking conditions cause variable losses (Lešková *et al*., 2006). Folate losses due to baking are quite common and reported by different scholars (Tamene *et al*., 2019; Lešková *et al*., 2006). However, more than 75 % of the folates were retained in some examples of baking *injera* (*ersho* fermented 100 % sorghum *injera* and *L. plantarum* fermented 100 % barley *injera*). This finding suggests that folate losses could be minimized if *injera* baking process optimized.

Based on portion size estimates obtained from the national food consumption survey, the average folate content in our *ersho* fermented *injera* samples made with different cereals (7.99 μg/100 g in tef and sorghum blended *injera*, 9.79 μg/100 g in wheat and sorghum blended *injera*, 8.94 μg/100 g in 100 % sorghum *injera* and 7.82 μg/100 g in 100 % barley *injera* per fresh weight) could contribute less than 5 % to the folate RNI of children (1–3 years) and woman of reproductive age. Use of the same cereals with equivalent proportions but changing the inoculum (*L. plantarum* instead of *ersho*) yields *injera* with better folate contents and contributed more than 5 % to the daily folate requirement of the two vulnerable population groups. Assuming two servings (∼350 g/serving) of *injera* are consumed per day by women of reproductive age, *injera* made with cereals other than tef could contribute more than10 % of the RNI if *L. plantarum* is used as inoculum. The present study clearly showed that all *injera* prepared using the traditional process had the lowest folate content as compared to its counterpart process which used *L. plantarum*.

The contribution of consumption of these *injeras* made using *L. plantarum* to folate requirement is a bit lower than tef *injera* fermented with the same strain (Tamene *et al*., 2022) and this could be obviously due to differences in the initial total folate content where significantly higher initial total folate is observed in tef flour (Tamene *et al*., 2019a) as compared to flours of any cereals used in this study. Otherwise, in both cases the strain *L. plantarum* is found to be effective in enhancing the folate content and makes the contribution twice as compared to contribution that could be obtained from consumption of *injera* made with the traditional process which uses *ersho* as inoculum.

Current studies have focused on folate improvement through fermentation with minimum emphasis to the organoleptic quality of the final product, which is one of the quality parameters for the food to be consumed. Our previous studies investigated the effect of *L. plantarum* on the sensory parameters of tef *injera* and we reported that all of the panellists accepted the folate enhanced tef *injera* made with the strain (Tamene *et al*., 2022). Similarly, in the present study, almost all *injera* samples fermented with *L. plantarum* was accepted by the panellists with the exception of 100 % sorghum *injera*, which requires further investigation to make it more acceptable as it has comparatively higher initial folate as compared to other cereals considered in the present study.

Certain limitations need to be considered in the interpretation of our findings. To what extent discarding of the supernatants contributed to folate loss was not quantified which would have enabled as to be clear with the cause of folate losses during fermentation (material loss or loss due to consumption by microorganisms). The study used a fixed proportion during blending of cereals but multiple proportion could yield a different result both on the folate content and sensory attributes of the final product. The present study focused only on few common cereals as there could be other cereals that can be used for making *injera* in other parts of the country.

## 5. Conclusion

Cereals such as wheat, barley, sorghum and tef could be considered as sources of folate and fermenting it to prepare *injera* can either increase or decrease its folate content, while baking always leads to folate losses. Either reduction or increment of folate contents could be observed in dough samples when cereals are fermented naturally using *ersho*. In case of fermentation which used *L. plantarum*, there was always increment of folate contents in all of the doughs. Though cooking or baking invariably leads to reduction of folate content in the final ready to be consumed staple food (*injera*), still those *injeras* made with different cereals and fermented with our strain (*L. plantarum*) had much better folate content as compared with same *injera* fermented with *ersho*. Accordingly, consumption of any of *injeras* fermented with *L. plantarum* contributed much more to folate RNI of children and women of reproductive ages than consumption of *injera* fermented with *ersho*. Though the folate content of *L. plantarum* fermented *injera* made with sorghum was better and contributed higher to folate RNI of both population groups, it was the least accepted product by the potential consumers. Our study has clearly indicated that *L. plantarum* could be used for the development of fermented foods like *injera* prepared with different cereal types with better folate content and consumer preferences. This research also showed that *L. plantarum* strain can be used as a potential folate source not only for *injera* made with tef but also for any *injera* that could be made with other cereals. This could open the door to novel food formulation with better folate content.

## Author Contributions

Conceptualisation, K.B. and A.T.; methodology, K.B. and A.T., validation, A.T. and K.B.; formal analysis, T.M. and A.T.; investigation, T.M. and A.T.; resources, K.B. and A.T.; writing—original draft preparation, T.M. and A.T.; writing—review and editing, K.B. and A.T.; visualisation, A.T. and K.B.; supervision, K.B. and A.T.; project administration, K.B.; funding acquisition, K.B. and T.M.

## Acknowledgments

Financial support was obtained from the graduate programme of Addis Ababa University. The authors would like to thank the panellists who involved during sensory analysis of the product. The authors also thank Ethiopian Public Health Institute for allowing us access to the National Food Science and Nutrition Laboratory for some of the laboratory works.

## Conflicts of Interest

The authors declare no conflict of interest.

## Notes

### Competing Interest Statement

The authors have declared no competing interest.

